# A novel histone deacetylase inhibitor-based approach to eliminate microglia and retain astrocyte properties in glial cell culture

**DOI:** 10.1101/2021.11.09.467827

**Authors:** Xi-Biao He, Yi Wu, Haozhi Huang, Fang Guo

**Affiliations:** Laboratory of Stem Cell Biology and Epigenetics, School of Basic Medical Sciences, Shanghai University of Medicine and Health Sciences, Shanghai 201318, China; Speech Therapy Department, The Second Rehabilitation Hospital of Shanghai, Shanghai 200441, China; Department of Orthopaedic Surgery, Shanghai Tenth People’s Hospital Affiliated to Tongji University, Shanghai 200072, China; Shanghai University of Medicine and Health Sciences Affiliated Zhoupu Hospital, Shanghai 201318, China

**Keywords:** astrocyte, microglia, HDAC inhibitor, inflammation, SARS-CoV-2

## Abstract

The close association between astrocytes and microglia causes great difficulties to distinguish their individual roles in innate immune responses in central nervous system. Current chemical-based methods to eliminate microglia in glial cell culture introduce various molecular and functional alterations to astrocytes. Here, we describe a novel two-step approach to achieve a complete elimination of microglia without affecting the biological properties of co-cultured astrocytes by temporal treatment of histone deacetylase inhibitor trichostatin A (TSA). We verify TSA as a potent inducer for microglial-specific cell death, which also causes comprehensive gene expression changes in astrocytes. However, withdrawal of TSA not only ensures no microglia repopulation, but also restores all the gene expression changes in terms of astrocyte functions, including neurotrophic factors, glutamate and potassium transporters, and reactive astrocyte subtypes. By contrast, withdrawal of PLX5622, the commonly used colony-stimulating factor 1 receptor inhibitor neither prevents microglia repopulation nor restores the gene expression changes mentioned above. Using this method, we are able to discriminate differential roles of microglia and astrocytes in the induced expression of antiviral and pro-inflammatory cytokines upon various pathological stimuli including the spike protein of SARS-CoV-2. This simple and efficient method can be customized for the understanding of microglia-astrocyte interaction and the development of epigenetic therapies that target over-activated microglia in neuroinflammation-related diseases.

## Introduction

Microglia and astrocyte are two major types of immune responding cells in the central nervous system. How the inductions of a variety of immune responses correspond to each cell type is not fully understood. The limitation is largely due to the close physical and chemical interactions between microglia and astrocytes in brain and in cell culture. In order to dissect their individual contribution in immune responses, it is necessary to separate microglia and astrocyte from each other before they are challenged by pathological stimuli. For in vivo studies, one of the most commonly used methods to eliminate resident microglia is by administration of colony-stimulating factor 1 receptor (CSF1R) inhibitors, which selectively affects microglial survival (Elmore et al., 2014). However, withdrawal of CSF1R inhibitors allows in situ microglial repopulation in brain (Huang et al., 2018). A complete elimination of resident microglia in brain is likely not preferred, as they might play detrimental or beneficial roles in different pathological conditions and neural disorders (Neumann et al., 2006, 2008; Lalancette-Hebert et al., 2007; Dagher et al., 2015; Acharya et al., 2016; Rubino et al., 2018; Spangenberg et al., 2019; Li et al., 2021), whereas for in vitro studies using mixed glial cell culture, a complete elimination of microglia is necessary to exclude all the physical and chemical contaminations derived from microglia that affect co-cultured cell types. A pure microglia-free cell niche provides unique advantages over mixed glial cell culture for the understandings of cell type-specific functions of these cells. Multiple drug-based approaches to eliminate microglia from astrocytes have been described, including CSF1R inhibitors (Giulian et al., 1993; Hamby et al., 2006; Kumamaru et al., 2012; Vilalta and Brown, 2014; Sepulveda-Diaz et al., 2016; Green et al., 2020). Despite the high efficiency of microglial elimination, one common potential disadvantage of these methods is that the chemicals used could most likely affect the biochemical characteristics of remaining astrocytes in an irreversible way, resulting in misleading interpretations for the subsequent experiments and analyses.

Antiviral and pro-inflammatory responses are two critical functions of glial cells for host innate immunity. Upon viral infection, a series of antiviral and pro-inflammatory cytokines were rapidly induced in microglia and astrocytes. Type I interferons (IFNs) such as IFNα and IFNβ represent one of the major mediators for antiviral response, leading to expression of hundreds of IFN-stimulated genes as effectors for viral defense, whereas pro-inflammatory response is composed of induced expressions of a wide range of cytokines and chemokines such as interleukin 6, interleukin 1β, tumor necrosis factor α and induced nitric oxide synthase. Microglia are generally regarded as the earliest responding cells of inflammation in the brain, which also modulates the subsequent activation of pro-inflammatory response in astrocytes. Under certain conditions, depletion of microglia by CSF1R inhibitor in vivo and in vitro attenuated the expression of pro-inflammatory molecules and resulted in beneficial outcomes (Feng et al., 2017; Coleman et al., 2020; Li et al., 2021). In other cases, experimental ablation of microglia exacerbated inflammation and worsen neural injuries (Lalancette-Hebert et al., 2007; Rubino et al., 2018). On the other hand, astrocytes have been reported as the main cell type for IFNβ production upon virus infection in vivo (Lindqvist et al., 2016; Pfefferkorn et al., 2016; Chhatbar et al., 2018). However, due to the complexity of the interaction between microglia and astrocytes, it would be very difficult to identify the exact cell type from which the immune responses were originated if the microglia and astrocytes were not completely separated from each other.

Histone acetylation is one common histone modification that dynamically affects chromatin structure and gene expression. The deacetylation of histone is catalyzed by a wide range of histone deacetylases and can be inhibited by a group of epigenetic drugs called HDAC inhibitors (HDACis). The specificity of HDACis varies. Trichostatin A (TSA) and Vorinostat (SAHA) are hydroxamates that non-specifically inhibit activity of both class I and class II histone deacetylases thus are regarded as pan HDACis. Valproic acid (VPA) and sodium butyrate, on the other hand, are short chain fatty acids that more specifically inhibit class I HDAC activity, whereas Tubastatin A (TubA) is a selective inhibitor of HDAC6. One of the most promising advantages of HDACis in clinical applications is that the epigenetic alterations induced are generally reversible with drug cessation (Ho et al., 2020). It has been reported that HDACis including TSA, VPA and sodium butyrate can induce microglia apoptosis (Chen et al., 2007), supporting the potential application of HDACis as eliminator of microglia from astrocytes. However, the specificity of HDACis to microglia and their impact on co-cultured astrocytes remain to be elucidated.

Here we describe a novel HDACi-based approach for the elimination of virtually all microglia from co-cultured astrocytes by specifically inducing microglia cell death. Withdrawal of HDACi after microglial elimination not only allowed no repopulation of microglia, but also brought back the epigenetic and transcriptional status of the remaining astrocytes to the levels before HDAC inhibition. This provides us an appropriate opportunity to determine how astrocytes, with or without microglial participation, respond to various immune stimuli including the spike protein of SARS-CoV-2.

## Materials and Methods

### Cell culture

Mixed glial cells were extracted from postnatal day 1 mouse brain cortices, mechanically dissociated and grown in Dulbecco’s modified Eagle medium (DMEM; Invitrogen) supplemented with 10% fetal bovine serum (Gibco) and 1% Penecillin/Streptomycin for three to four weeks until full confluency. Cells were passaged using mild trypsin method (Saura et al., 2003) with some modifications. In brief, cells were washed by phosphate buffer saline (PBS) twice and incubated in a mixture of 0.25% trypsin-EDTA and DMEM at a ratio of 1:4 for 40 minutes until the upper layer which consisted of astrocytes and microglia completely detached. The upper layer was removed, mechanically dissociated and plated for mixed glial cell experiments, and the remaining adherent layer was enzymatically dissociated with 0.25% trypsin-EDTA and plated for pure microglia experiments. The mixed glial cells were cultured for 5-7 days until a flat monolayer formed. Pure microglial cells were further cultured for three to five days to make sure that the majority of cells recovered and remained in a ramified morphology. For drug treatments, following inhibitors were used: 1 μM TSA, 200 nM SAHA, 4 mM VPA, 40 μM TubA (all from Sigma) and 20 μM PLX5622 (PLX; Selleck). Drugs were added with medium change once every three days. For drug withdrawal experiments, cells were washed with PBS twice after drug-containing media were removed, then replaced with fresh drug-free media. Cells were incubated in 5% CO2, 37°C incubator.

### Pathological stimulation

For SARS-CoV-2 pseudovirus generation, the plasmid carrying spike protein sequence (Miaoling Plasmid Sharing Platform plasmid #P18156) was co-transfected with psPAX2 (Addgene plasmid #12260) into HEK293T cell line using Lipofectamine 2000 (Invitrogen) according to manufactor’s instruction for 6 hours and cells were replaced with fresh media containing DMEM supplemented with 10% fetal bovine serum. 24 hours later, supernatants were harvested, filtered and stored at - 80°C until use. For cell stimulation, half glial cell culture media were replaced with same amount of virus supernatants for 6 hours. To confirm the specificity of the spike protein, an inhibitory antibody of the spike protein (SinoBiological) was 1-hour pre-treated before virus supernatants. For lipopolysaccharide (LPS) and virus surrogate poly (I:C) stimulation, 200 ng/mL LPS and 20 μg/mL Poly (I:C) were directly added to fresh cell culture media for 6 hours.

### Lactate dehydrogenase (LDH) assay

The LDH assay was performed according to manufactor’s instructions (Beyotime) to measure the cytotoxicity in glial cells. Cytotoxicity was measured by subtracting LDH content in HDACi-treated cells from total LDH in untreated controls.

### Immunofluorescence staining and imaging

Cells were fixed with 4% paraformaldehyde for 20 minutes, permeabilized by 0.3% Triton-X100 and blocked by 1% bovine serum albumin for 40 minutes, then incubated with primary antibodies at 4□ overnight. Alexa Fluo series of second antibodies (Thermo Scientific) were applied accordingly for one hour at room temperature. Cells were finally mounted in 4’,6-diamidino-2-phenylindole (DAPI) and examined using fluorescence microscope (Leica DMi8). The first antibodies include rabbit anti-Iba1 (Wako), mouse anti-GFAP (Sigma) and rabbit anti-Ki67 (Abcam).

### Western blot

Histones were acid-extracted from cell samples. 3 µg of histone samples were electrophoresized with 15% SDS-PAGE gels, and the blotted membranes were incubated with anti-histone 3 acetylation(H3Ac), histone 3 lysine 27 tri-methylation (H3K27me3) and histone 3 antibodies (all from Millipore). Positive bands were detected by chemiluminescent reagents (Thermofisher) and quantified using Image J software. Quantification was performed using average values from three independent experiments.

### Real-time PCR analysis

RNA from cell samples was extracted, reverse-transcribed (TaKaRa), amplified, and applied to real-time PCR analyses (Roche). The comparative cycle threshold method was used for quantification. All the experiments have been independently repeated two to three times from different batches of cell cultures to guarantee same trends of gene expression. Data from one representative experiment were used to generate each histogram. Error bar represents standard derivation from three replicates of one single experiment. The primer sequences are listed in Supplemental Table 1. The heatmaps were generated using Graphpad Prism based on real-time PCR results using the mean values from three replicates of one single experiment in a form of log2 fold change. Detailed quantification data can be found in Supplemental Table 2.

### Cell counting and statistics

Immunoreactive or DAPI-stained cells were counted in at least 10 random regions of each culture coverslip using an eyepiece grid at a magnification of 200X. Data are expressed as mean ± S.E.M. of three independent cultures. Statistical comparisons were made using Student’s t-test or one-way ANOVA with Tukey’s post hoc analysis (Graphpad Prism).

## Results

### HDAC inhibitors induce microglia-specific cell death

To determine the effect of different HDAC inhibitors on the survival of microglia and astrocytes, heterogeneous cultures mainly composed of astrocytes and microglia extracted from postnatal day 1 mouse cortices were challenged with pan HDAC inhibitors TSA and SAHA, class IIa HDAC inhibitor VPA and class IIb HDAC inhibitor TubA respectively for two days. In parallel, CSF1R inhibitor PLX was treated as a well-defined positive control. Remarkable loss of ionized calcium binding adaptor molecule 1 (Iba1) positive microglia was induced by 1 μM TSA, 200 nM SAHA, 40 μM TubA, and to a less extent by 4 mM VPA and 20 μM PLX (Fig. 1A-B), indicating that these inhibitors affected microglial survival rather than proliferation. Nearly-complete elimination of Iba1+ microglia was achieved by extending the treatment periods of PLX and VPA to three or five days, respectively (Fig. 1C), suggesting that the efficiency of different HDACis on microglial survival varied. By contrast, none of the inhibitors significantly affected the number of glial fibrillary acidic protein (GFAP) positive astrocytes (Fig. 1A and 1D). Consistently, LDH assay and Ki67 staining confirmed that none of these HDACis affected the survival or proliferation of astrocytes (Supplemental Fig. 1A-B). In addition, gene expression analysis demonstrated dramatic decrease of microglia-specific gene transcripts including *Csf1r, Spi1* or *Dap12* after 5 days of PLX or HDACi treatments (Fig. 1E). These results demonstrate a microglia-specific viability response to the inhibition of HDACs.

**Fig. 1.**
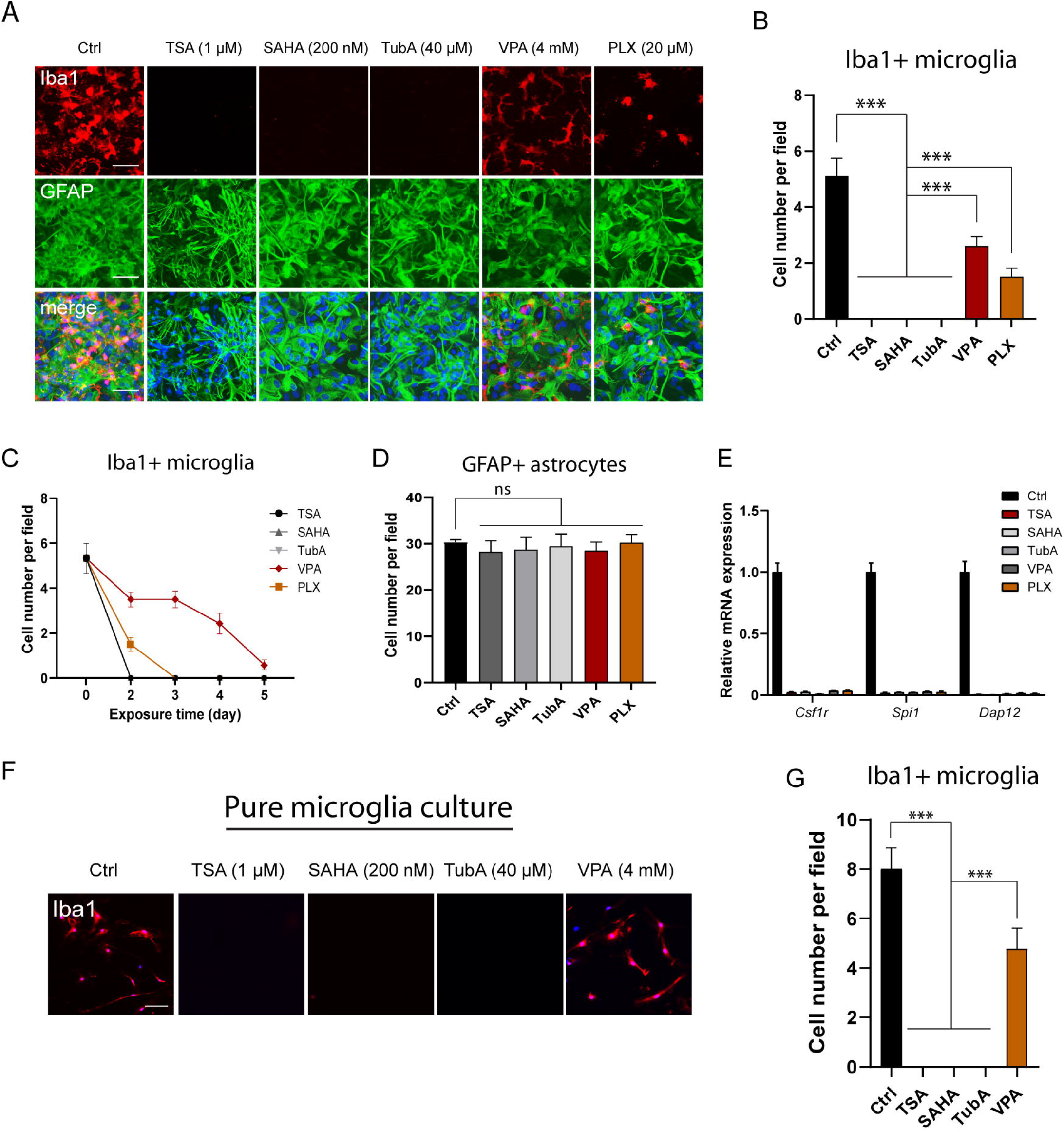
Different HDAC inhibitors affected microglial survival. (A) Representative immunofluorescence images showing the effect of different HDAC inhibitors and CSF1R inhibitor on the survival of co-cultured mouse microglia and astrocytes after two days of drug exposure. Pan HDAC inhibitors Trichostatin A (TSA; 1μM) and Vorinostat (SAHA; 200 nM) and class IIb HDAC inhibitor Tubastatin A (TubA; 40 μM) induced extensive dose-dependent loss of Iba1+ microglia, and to a less extent by class IIa HDAC inhibitor valproic acid (VPA; 4 mM) and CSF1R inhibitor PLX5622 (PLX; 20 μM), while GFAP+ astrocytes were not affected. Cell nuclei were counterstained by DAPI. (B) The quantification of microglial cell number with treatment of different inhibitors for 2 days on co-cultured microglia and astrocytes. (C) Cell number of Iba1+ microglia in response to treatment of different inhibitors for 5 days on co-cultured microglia and astrocytes. (D) Cell number of GFAP+ astrocytes with treatment of different inhibitors for 2 days on co-cultured microglia and astrocytes. (E) Real-time PCR analysis of microglia-specific gene transcript expression after treatment of PLX or HDAC inhibitors for 5 days on co-cultured microglia and astrocytes. (F) Representative immunofluorescence images showing the effect of different HDAC inhibitors (same concentrations as in co-culture) on the survival of pure mouse microglia. Cells were treated for 2 days. (G) The quantification of microglial cell number in pure microglia culture. Scale bars represent 20 μm. Cell numbers were counted in 10 random areas of each culture coverslip. Data represent mean ± S.E.M. n=3 independent culture. *** P < 0.001; ns, not significant; one-way ANOVA with Tukey’s post hoc analysis.

Astrocyte-derived factors such as CSF1, TGF-β2 and cholesterol have been shown to be essential for microglia survival (Bohlen et al., 2017). To determine whether HDACi-induced microglial cell death is dependent on astrocytes, we applied same treatments of HDACis to a homogeneous culture of microglia, which was derived from the heterogeneous glial culture by a specific passaging method (Saura et al., 2003). Similar results of cell loss were observed by 2 days treatments of TSA, SAHA and TubA, whereas effect of VPA was much milder (Fig. 1F-G). However, the loss of microglia in homogenous microglia culture within 2 days of treatments appeared to be more predominant than that of astrocyte-microglia co-culture, likely reflecting a supportive function of astrocytes for microglia survival. Nevertheless, these results demonstrate that HDAC inhibition is specifically detrimental to microglia and could serve as an efficient approach for microglial elimination in glial cells.

### No microglia repopulation by withdrawal of TSA

Repopulation of microglia has been reported in vivo after treatment of CSF1R inhibitor, likely from maturation of microglia progenitors (Elmore et al., 2014; Zhan et al., 2020) and/or rapid self-renewal of surviving microglia (Huang et al., 2018). To determine whether this would also be the case for HDACis, glial cells were treated with TSA, SAHA, TubA or PLX for 3 days and re-incubated in inhibitor-free culture media for as long as a week. As expected, withdrawal of PLX resulted in repopulation of Iba1+ microglia. However, no Iba1+ cells were found after withdrawal of TSA or SAHA. Interestingly, withdrawal of TubA caused emergence of some microglial cells (Fig. 2A-B), suggesting a diversity of the effect and likely the mode of action of different HDACis on microglial survival. In addition, long-term treatment of TubA caused cell death of GFAP+ astrocytes (Supplemental Fig. 2). At this point, we have optimized cell culture conditions and selected pan HDACi TSA for the following experiments to ensure a complete elimination of microglia and/or their progenitors with minimal impact on the cell number of astrocytes (illustrated in Fig. 2E).

**Fig. 2.**
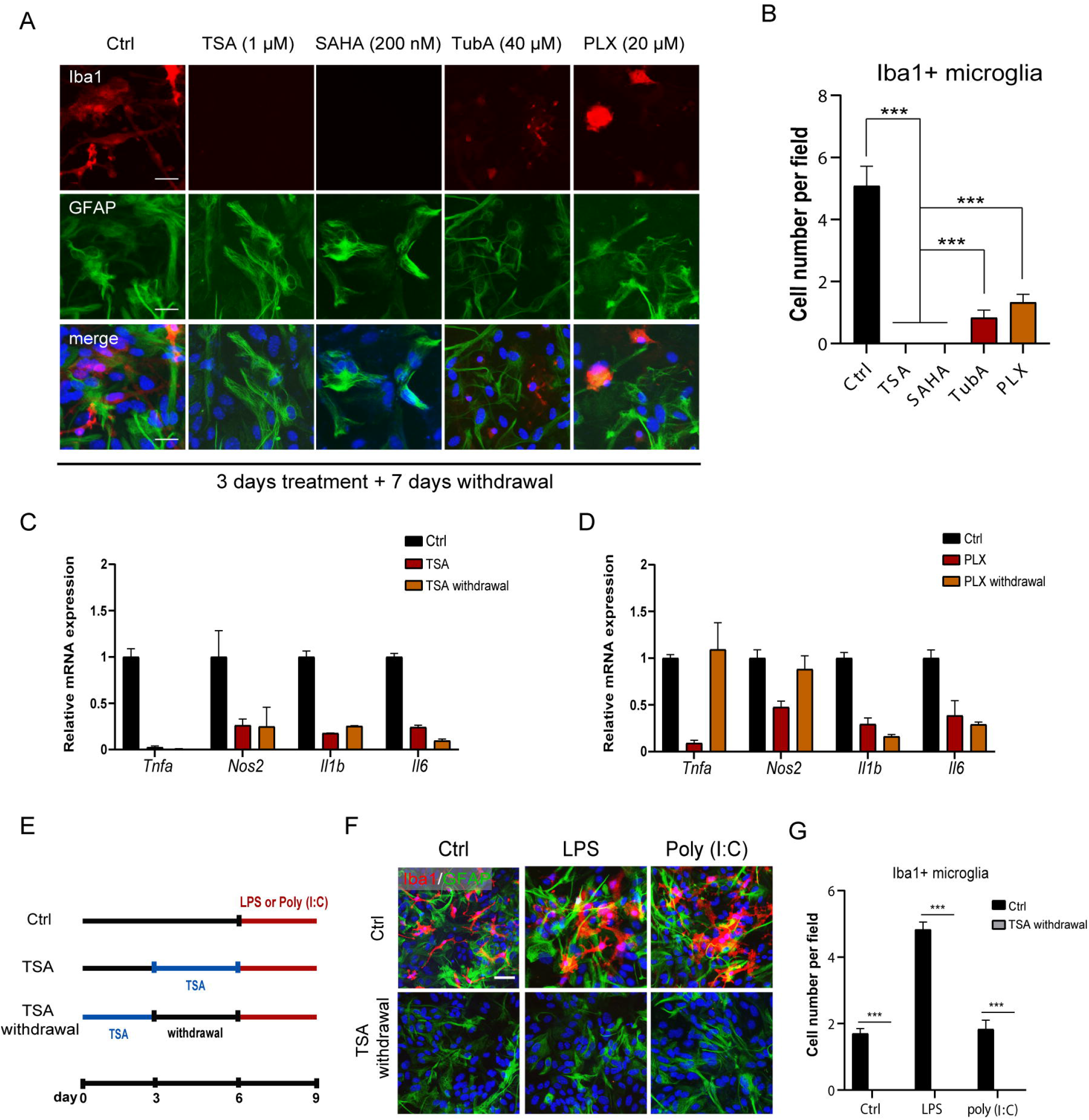
Withdrawal of HDAC inhibitors and CSF1R inhibitor on microglia repopulation. (A) Representative immunofluorescence images showing co-cultured Iba1+ microglia and GFAP+ astrocytes treated with different HDAC inhibitors including Trichostatin A (TSA), Vorinostat (SAHA) and Tubastatin A (TubA) and CSF1R inhibitor PLX5622 (PLX) for 3 days and then withdrawn of inhibitors for 7 days. Cell nuclei were counterstained by DAPI. (B) The quantification of microglial cell number after sequential treatments and withdrawals of various inhibitors. (C) Real-time PCR analysis of pro-inflammatory cytokine gene transcripts (Tnfa, Il1b, Il6 and Nos2) after treatment and withdrawal of TSA in co-cultured microglia and astrocytes. (D) Real-time PCR analysis of same four pro-inflammatory cytokines gene transcripts after treatment and withdrawal of PLX in co-cultured microglia and astrocytes. (E) Schematic illustrating the procedure of pathogen stimulation including bacterial lipopolysaccharide (LPS) and virus surrogate poly (I:C). The TSA withdrawal procedure without pathogen stimulation is the standard approach for microglia elimination. (F) Representative immunofluorescence images showing co-cultured Iba1+ microglia and GFAP+ astrocytes after treatment and withdrawal of TSA and pathogen stimulation. (G) The quantification of Iba1+ cell number after pathogen stimulation. Scale bars represent 20 μm. Cell numbers were counted in 10 random areas of each culture coverslip. Data represent mean ± S.E.M. n=3 independent culture. *** P < 0.001; ns, not significant; one-way ANOVA with Tukey’s post hoc analysis.

Pro-inflammatory cytokines are constitutively expressed at basal level in astrocytes and microglia under normal culture condition. To establish a direct link between inflammation induction and the presence of microglia, we investigated the consequences of treatment and withdrawal of TSA and PLX on the transcription of four pro-inflammatory cytokine genes such as *Tnfa, Il1b, Il6* and *Nos2*. Treatments of TSA and PLX in astrocyte-microglia co-culture remarkably reduced the overall mRNA levels of these genes, suggesting that their expressions were overwhelmingly contributed by microglia. The expression levels of these genes were not altered after withdrawal of TSA (Fig. 2C), confirming on the molecular level that microglia repopulation did not occur. This result also reflected that treatment of TSA did not regulate pro-inflammatory cytokine gene expression in microglia-free astrocytes. By contrast, withdrawal of PLX up-regulated the expressions of *Tnfa* and *Nos2*, but not *Il1b* and *Il6* (Fig. 2D), which were likely to be derived from repopulated microglia. These results confirmed the consequent differences between withdrawal of TSA and PLX in terms of microglia repopulation and the basic level of inflammation in astrocytes.

Pathological stimuli are strong inducers for microglial self-renewal and repopulation. To further validate the complete elimination of microglia by TSA, glial cells that underwent 3-day sequential treatment and withdrawal of TSA were challenged with bacterial LPS or virus surrogate poly (I:C) for another 3 days (Fig. 2E). As expected, neither stimulus triggered any emergence of Iba1+ cells (Fig. 2F-G). Taken together, these results demonstrate that temporal pan HDAC inhibition serves as an efficient means to eliminate virtually all the microglia and/or their progenitors through an alternative pathway of CSF1R inhibition.

### Molecular signatures of astrocytes are restored by withdrawal of TSA but not PLX5622

Epigenetic modifications are commonly reversible after removal of the inducers (Allis and Jenuwein, 2016). As PLX and TSA seem not affecting the survival and proliferation of astrocytes, we sought to address whether the expressions of astrocyte characteristic genes were affected by these inhibitors, and whether withdrawal of TSA or PLX could restore those changes. Glial cell culture was exposed to TSA or PLX for 3 days for complete elimination of microglia, and the remaining astrocytes were incubated in replaced media without inhibitors for another 3 days in attempt to allow epigenetic recovery. Indeed, the global level of H3Ac was increased by TSA at day 3 and recovered at day 6 by the withdrawal of TSA. Interestingly, the level of H3K27me3 was decreased by TSA and also recovered after TSA withdrawal, suggesting a crosstalk between histone acetylation and histone methylation (Fig. 3A).

**Fig. 3.**
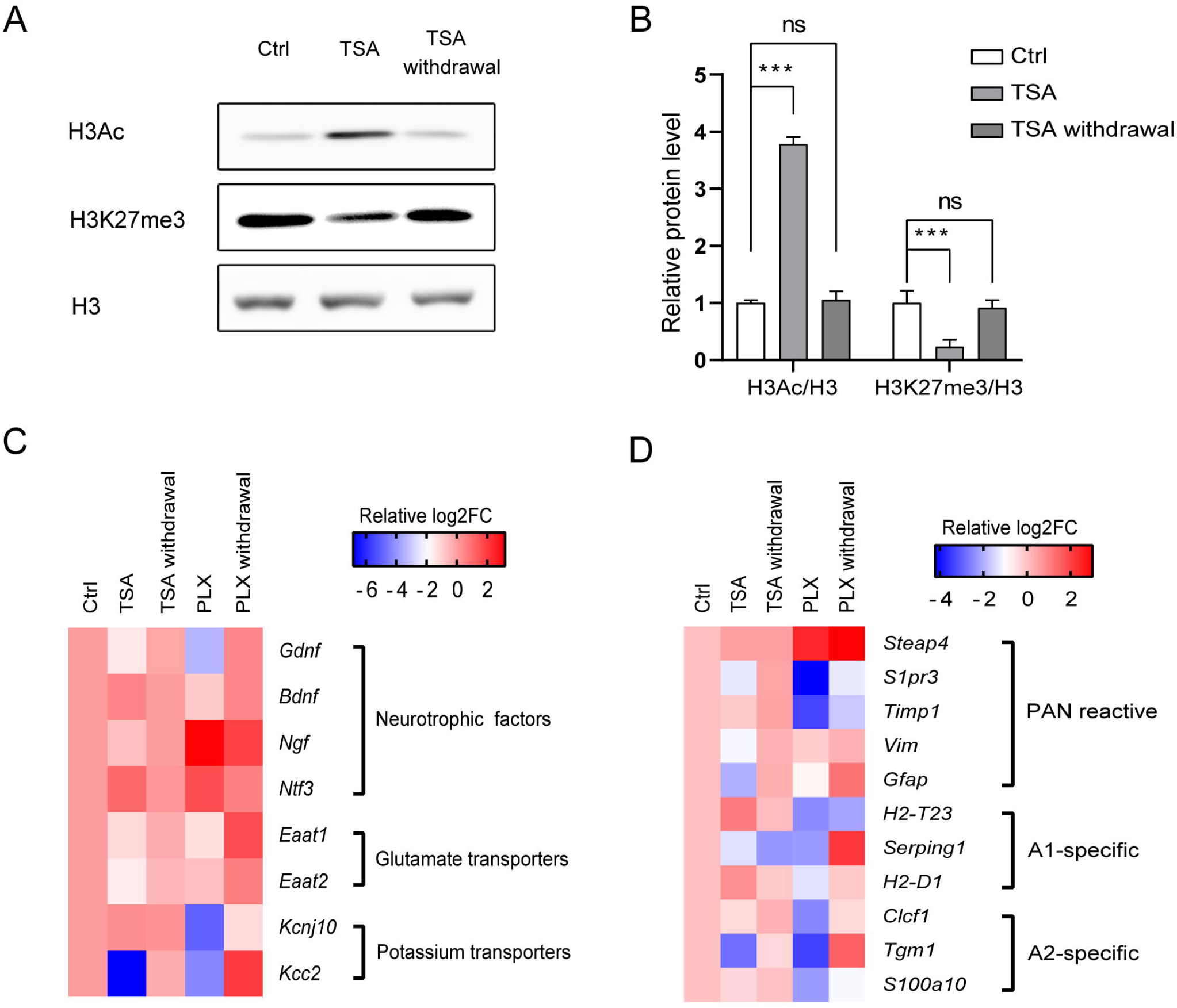
Epigenetic status and astrocyte molecular signatures are reversible by withdrawal of HDAC inhibitor. (A) Representative immunoblot images of histone modifications in glial cells treated with Trichostatin A (TSA) and first treated then withdrawn with TSA. Expression of histone 3 acetylation (H3Ac) and histone 3 lysine 27 tri-methylation (H3K27me3) were shown and histone 3 (H3) was used as reference. (B) The quantification of H3Ac and H3K27me3 in reference of H3. mean ± S.E.M. n=3 independent culture. *** P < 0.001; ns, not significant; one-way ANOVA with Tukey’s post hoc analysis. (C) Heat map showing expression changes of astrocyte-related genes after treatments and withdrawals of TSA and CSF1R inhibitor PLX5622 (PLX). Expression level is scaled as relative log2FC (fold change). (D) Heat map showing expression changes of astrocyte subtype genes after treatments and withdrawals of TSA and PLX. Expression level is scaled as relative log2FC.

To determine how astrocytes were dynamically affected during the process of microglia elimination, gene expression analysis was performed to monitor the transcriptional changes of a wide range of genes that were extensively involved in astrocyte biology. Comprehensive transcriptional changes of genes specific to astrocyte functions were induced by treatments of TSA and PLX, including neurotrophic factor genes *Gdnf, Bdnf, Ngf* and *Ntf3*, glutamate transporter genes *Eaat1* and *Eaat2*, and potassium transporter genes *Kcnj10* and *Kcc2*. Interestingly, withdrawal of TSA significantly restored all of them to the original levels in microglia-astrocyte co-culture. On the contrary, fewer genes were restored by withdrawal of PLX (Fig. 3B). These results suggest that astrocytes underwent treatment and withdrawal of TSA share more molecular signatures with co-cultured ones.

Recent studies have classified two reactive subtypes of astrocytes, each of which corresponds to the up-regulation of a core set of subtype-specific genes (Liddelow et al., 2017; Clarke et al., 2018). We further investigated how these genes were affected by treatments and withdrawals of TSA and PLX. Amongst all the genes examined, five were classified as pan reactive, three as A1 specific, and three as A2 specific. Interestingly, although both treatments of TSA and PLX caused the transcriptional alterations of all these genes, the trends were not consistent with any of the subtype criteria, suggesting that unique molecular patterns were induced by TSA and PLX. However, withdrawal of TSA restored almost all of them to the original levels in co-culture, in contrast to the substantial diversity from withdrawal of PLX (Fig. 3C). Taken together, these results suggest that the application of HDACi has an advantage over CSF1R inhibitor in retaining astrocyte properties after microglia are eliminated.

### Elimination of microglia blunts inflammatory but not antiviral response in astrocytes

The implication of SARS-CoV-2 in central nervous system has raised increasing concerns. The spike protein of SARS-CoV-2 has been reported to cause inflammation in macrophage (Shirato and Kizaki, 2021). We took advantage of the findings above in order to compare the immune responses derived from astrocyte-microglia co-culture and astrocyte-only culture from various pathological stimuli including LPS, Poly (I:C) and the spike protein of SARS-CoV-2. To do so, co-cultured astrocytes and microglia without TSA treatment and after treatment and withdrawal of TSA were challenged with LPS, Poly (I:C) and spike protein-enveloped pseudovirus supernatant for 6 hours. All three stimuli significantly induced the transcription of four pro-inflammatory cytokines in microglia-astrocyte co-culture. By contrast, their expression levels were remarkably lower in astrocyte-only culture, suggesting blunted activation of these genes without involvement of microglia (Fig. 4A). The specificity of the spike protein was confirmed by treating cells with an inhibitory antibody (Supplemental Fig. 3). These results confirm the central role of microglia in initiation of the pro-inflammatory response and suggest similarities of pro-inflammatory response among SARS-COV-2, LPS and Poly (I:C) in microglia and astrocytes.

**Fig. 4.**
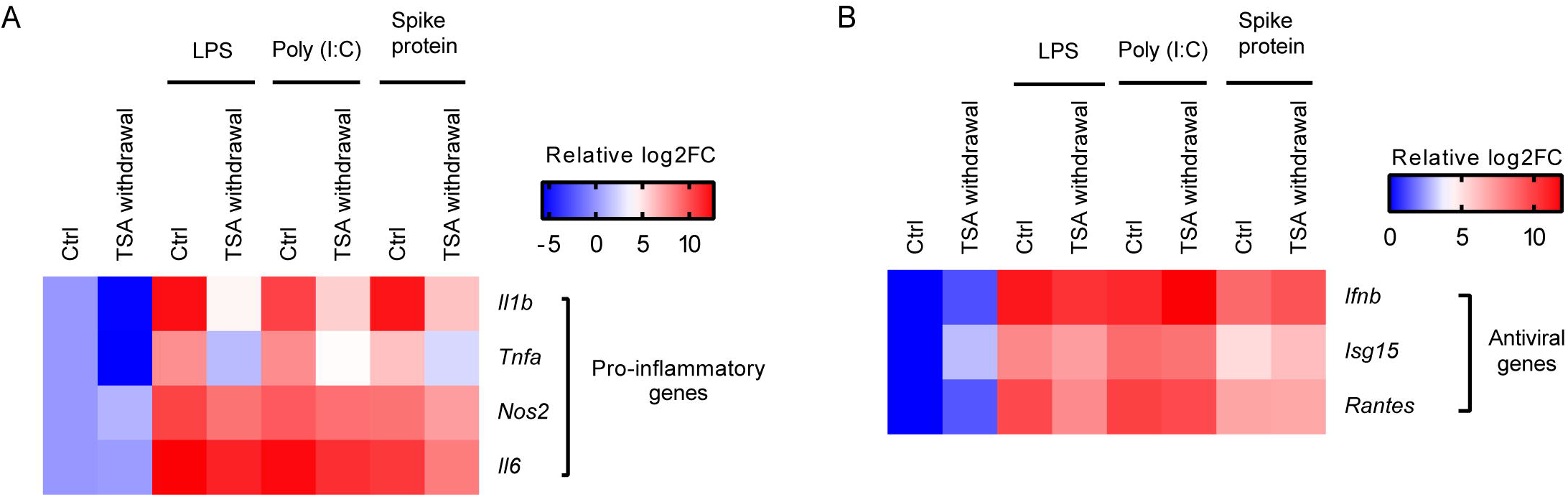
Pro-inflammatory and antiviral responses to various pathological stimuli in astrocytes with or without microglia. The astrocyte-only culture was derived from microglia-astrocyte co-culture by treatment of Trichostatin A (TSA) for 3 days and withdrawal of TSA for another 3 days. The astrocyte-only culture and microglia-astrocyte co-culture were then challenged for 6 hours by pathological stimuli including bacterial lipopolisaccharide (LPS), virus surrogate Poly (I:C) and the pseudovirus supernatant enveloped by the spike protein of SARS-CoV-2. Heat maps showing expression changes of four pro-inflammatory cytokine genes and three antiviral genes are shown in (A) and (B), respectively. Expression level is scaled as relative log2FC (fold change).

IFN-dependent antiviral genes encode for another important group of cytokines specifically induced for host immune defense against pathogenic viruses. In microglia-astrocyte co-culture, antiviral genes such as *Ifnb, Isg15* and *Rantes* were dramatically up-regulated in response to LPS, Poly (I:C) and spike protein stimulations, suggesting that strong antiviral response could be activated by these pathogens in microglia and/or astrocytes. However, different as pro-inflammatory cytokine genes, these genes were either slightly down-regulated upon LPS, or not altered upon poly (I:C) and the spike protein in astrocyte-only culture (Fig. 4B), indicating that microglia was either not involved or indispensable in comparison to astrocytes in the antiviral response. Collectively, these findings suggested differential roles of microglia and astrocytes in pro-inflammatory and antiviral responses to pathological stimuli, including SARS-CoV-2.

## Discussion

Primary culture of glial cells is commonly contaminated with certain amount of microglia. Even a minimal contamination of microglia is able to initiate amplifying immune responses in astrocytes (Saijo et al., 2009), introducing difficulties to study the microglia-independent characteristics of astrocytes. Concerns might be raised on usage of CSF1R inhibitor, a potent eliminator for certain amount of microglial population but not other CSF1R inhibition-resistant progenitors (Zhan et al., 2020), to provide a bona fide microglia-free context especially when the drug application is discontinued. Indeed, we verified that repopulation of Iba1+ microglia occurred after withdrawal of PLX (Fig. 2A-B). The re-emergence of microglia represents an incomplete purification of astrocytes which might lead to misinterpretation of experiments using PLX-treated cells as microglia-free astrocytes. For instance, PLX withdrawal caused remarkable inflammatory activation in response to LPS or Poly (I:C) in comparison to nearly non-responsiveness in PLX-treated cells (Fig. 2D). This activation was likely derived from repopulated microglia, rather than effects induced by PLX withdrawal or remaining astrocytes. However, there was no microglia repopulation after TSA treatment or TSA withdrawal, demonstrating a wider range of, if not all, microglia population responding to HDAC inhibition than CSF1R inhibition.

The main aim of this study is to provide an easy and customizable approach to separate astrocytes from microglia in mixed glial cell culture. A key question following our in vitro results is whether this approach is also applicable in vivo. Unfortunately, we failed to see loss of Iba1+ cells in the hippocampal regions of two-month-old adult mice, after they had received 6 days of intraperitoneal injections of TSA and SAHA (unpublished data). Therefore, we conclude that the present approach is applied only in vitro to scientifically address differential roles for microglia and astrocytes. However, it would be too early to reach a definite conclusion that HDACis are not suitable for elimination of microglia in vivo, considering that the half-life of TSA and SAHA are only 2 hours after oral administration in human. It took at least two days of drug treatment to reach microglia elimination in our culture system. Perhaps a more sustained drug exposure given by other delivery methods such as diet chow in future study would help address this issue. On the other hand, a complete elimination of microglia in vivo might not be preferred for translational purposes, as accumulating evidence have demonstrated a Janus face of microglia in neurological disorders including stroke, trauma and neurodegenerative diseases (Neumann et al., 2006, 2008; Lalancette-Hebert et al., 2007; Dagher et al., 2015; Acharya et al., 2016; Rubino et al., 2018; Spangenberg et al., 2019; Li et al., 2021). Combining more experimental evidence from in vivo and in vitro studies using different approaches to eliminate microglia upon various pathological stimuli might help solve this controversy.

Mounting evidence have linked brain conditions in health and disease with diverse astrocyte reactivity, namely detrimental A1 and beneficial A2 subtypes. It is not surprising that treatments of TSA and PLX induced alterations of gene expression inconsistent with the molecular signature of either subtype, given that the category of these subtypes was derived from neuropathological conditions such as LPS and middle cerebral artery occlusion. The recovery of most of subtype-specific genes after TSA withdrawal confirmed that they were originally affected by TSA, which also suggested the potential role of histone acetylation in the transcriptional regulation of these genes. Interestingly, the gene expression changes from PLX withdrawal were different as those from either no treatment or PLX-treatment. As microglia repopulation was observed after PLX withdrawal, this result might reflect a previously unknown impact of repopulated microglia on the expression of those genes in astrocytes.

Latest research have demonstrated that SARS-CoV-2 can invade central nervous system and cause neurological and psychiatric outcomes in COVID-19 patients (Meinhardt et al., 2021; Taquet et al., 2021). Concerns have been raised on implication of SARS-CoV-2 for the inflammatory storm evoked by the virus in brain. The spike protein of SARS-CoV-2 has been shown to trigger immune responses in macrophage and monocytes (Shirato and Kizaki, 2021; Zhao et al., 2021). Multiple evidence exist confirming the expression of angiotensin-converting enzyme 2, the receptor for the spike protein of SARS-CoV-2 in astrocytes (Gowrisankar and Clark, 2016; Chen et al., 2021; Hernández et al., 2021). However, it remains largely unclear whether the spike protein should activate the same receptor recognition and signalings as bacterial LPS and double stranded RNA viral surrogate Poly (I:C) in glial cells, and to what extent do they differ. Therefore, we utilized a spike protein-enveloped pseudovirus supernatant to mimic the virus-cell interaction as a third pathogen stimulus besides LPS and Poly (I:C) and investigated how this interaction evoked pro-inflammatory and antiviral responses in microglia and astrocytes. The pro-inflammatory response was remarkably activated by spike protein in microglia-astrocyte co-culture and largely blunted in astrocyte-only culture. This result resembled the pro-inflammatory responses activated by LPS and Poly (I:C), suggesting that these stimuli might share same recognition receptors and/or merged signaling pathways of inflammatory response in microglia and astrocytes. Indeed, TLR4-dependent LPS recognition and TLR3-dependent Poly (I:C) recognition share merging downstream signaling pathways (Pitha, 2004) and recent study has reported that SARS-CoV-2 spike protein interacts and activates TLR4 to induce pro-inflammatory response (Zhao et al., 2021). The significant attenuation of expression levels of pro-inflammatory genes in astrocyte-only culture upon the spike protein also demonstrates a critical role of microglia in inflammatory response to SARS-CoV-2. On the other hand, the antiviral genes activated by the spike protein and Poly (I:C) were not attenuated but rather enhanced (for instance, *Ifnb*) in astrocyte-only culture. This surprising result suggests that different with inflammatory response, astrocytes might work independently from microglia and play major roles in antiviral response to RNA viruses such as SARS-CoV-2. However, observations from in vitro experiments do not necessarily reflect the in vivo reality. The culturing systems might change global microglia profile, especially their capability responding pathological stimuli. We could not exclude the possibility that the antiviral capacity is compromised in cultured microglia. Confirmation of this result using different culture systems and animal experiments are needed.

## List of Abbreviations

TSA: Trichostatin A
CSF1R: Colony stimulating factor 1 receptor
IFN: Interferon
HDAC: Histone deacetylase
HDACis: Histone deacetylase inhibitors
SAHA: Vorinostat
VPA: Valproic acid
TubA: Tubastatin A
PLX: PLX5622
LPS: lipopolysaccharide
DAPI: 4’,6-diamidino-2-phenylindole
Iba1: Ionized calcium binding adaptor molecule 1
GFAP: Glial fibrillary acidic protein
H3Ac: Histone 3 acetylation
H3K27me3: Histone 3 lysine 27 tri-methylation

## Declarations

### Ethics approval and consent to participate

All animal experiments were approved by the animal ethics committee of Shanghai University of Medicine & Health Sciences and have been performed in accordance with the ethical standards laid down in the 1964 Declaration of Helsinki and its later amendments.

### Consent for publication

Not applicable.

### Availability of data and materials

The data sets generated and analyzed during the current study are available from the corresponding author upon reasonable request.

### Authors’ contributions

XBH conceived the study, performed the experiments and wrote the manuscript. YW and HH was involved in analysis and interpretation of data, and manuscript drafting. FG was involved in experiment conduction. All authors have read and approved of the final version of the manuscript.

## Acknowledgements

We thank Jiaqing Yan, Kexuan Li, Kerun Chen and Yongjun Ma for technical help on immunostaining experiments.

---

**Supplemental Fig.1 Cytotoxicity and proliferation of astrocytes in astrocyte-microglia co-culture in response to HDAC inhibitors.** (A) Lactate dehydrogenase (LDH) assay was performed to measure cytotoxicity in astrocytes after 2 days treatment of various HDAC inhibitors. (B) Counting of percentage of GFAP+ astrocytes expressing proliferation marker Ki67 out of total GFAP+ cells after 2 days treatment of various HDAC inhibitors. Cell numbers were counted in 10 random areas of each culture coverslip. Data represent mean ± S.E.M. n=3 independent culture. ns, not significant; one-way ANOVA with Tukey’s post hoc analysis.

**Supplemental Fig.2 Long-term effect of HDAC inhibitors on astrocyte cell number in astrocyte-microglia co-culture.** Cells were incubated with HDAC inhibitors for 10 days, with drug and media replacement every three days. Cells were immunostained for GFAP (A) and counted (B). Cell numbers were counted in 10 random areas of each culture coverslip. Data represent mean ± S.E.M. n=3 independent culture. *** P < 0.001 compared with Ctrl; one-way ANOVA with Tukey’s post hoc analysis.

**Supplemental Fig. 3 Confirming the specificity of the spike protein-enveloped pseudovirus supernatants.** Astrocyte-microglia co-culture after standard TSA treatment and withdrawal procedure was challenged for 6 hours by the spike protein-enveloped virus supernatant. An inhibitory antibody against the spike protein was 1-hour pre-treated before virus supernatant. Gene expression changes of four pro-inflammatory cytokines and three antiviral genes were analyzed by real-time PCR experiments. Expression level is scaled as relative log2FC (fold change) and presented as heat map.

