## Supplemental Table 1 for "A novel histone deacetylase inhibitor-based approach to eliminate microglia and retain astrocyte properties in glial cell culture"

**Supplemental Table S1.** Oligonucleotide primers

| Primer name | Forward | Reverse |
| --- | --- | --- |
| Il6 | ATGGATGCTACCAAACTGGAT | TGAAGGACTCTGGCTTTGTCT |
| Tnfa | CGTCAGCCGATTTGCTATCT | CGGACTCCGCAAAGTCTAAG |
| Il1b | GCCCATCCTCTGTGACTCAT | AGGCCACAGGTATTTTGTCG |
| Nos2 | CCTCCTTTGCCTCTCACTCTTC | AGTATTAGAGCGGTGGCATGGT |
| Csf1r | ATGTGTGGTCCTACGGCATC | AGGCTGGGCCATTTGGTATC |
| Spi1 | TTTCTCCGCACACCATGTCC | AGGACGTGCATCTGTTCCAG |
| Dap12 | ACTGTGGTGTCCAGTGCATA | TCACGGAAGAACAGTCGCAT |
| Gdnf | TATTGCAGCGGTTCCTGTGA | TCCACACCGTTTAGCGGAAT |
| Bdnf | GTGACAGTATTAGCGAGTGGG | GGGTAGTTCGGCATTGC |
| Ngf | TTTGATCGGCGTACAGGCAG | TATTGGTGCAGTAGGGGCAC |
| Ntf3 | GGGTAGTTCGGCATTGC | GGCACACACACAGGAAGTGTC |
| Eaat1 | ACCAGGCGTTCTTAGGATGAC | TTTAAACCTGGGAGCTGCCTC |
| Eaat2 | AGTCAATGTGGTGGGCGATT | TCGTCGTAAATGGACTGCGT |
| Kcnj10 | TAAAAGATCTCCCGCTCCGC | AGCTTCTCGGGGTCTCCATA |
| Kcc2 | ACCGTTGTCTTTGTGGGTGT | ATCGGGAAATTGGGTGGGTC |
| Ifnb | CAGCTCCAAGAAAGGACGAAC | GGCAGTGTAACTCTTCTGCAT |
| Isg15 | CAGGACGGTCTTACCCTTTCC | AGGCTCGCTGCAGTTCTGTAC |
| Rantes | ATATGGCTCGGACACCACTC | ACTTGGCGGTTCCTTCGAG |
| Steap4 | CCCGAATCGTGTCTTTCCTA | GGCCTGAGTAATGGTTGCAT |
| S1pr3 | AAGCCTAGCGGGAGAGAAAC | TCAGGGAACAATTGGGAGAG |
| Timp1 | AGTGATTTCCCCGCCAACTC | GGGGCCATCATGGTATCTGC |
| Vim | TGGATCAGCTCACCAACGAC | AAGGTCAAGACGTGCCAGAG |
| Gfap | AGAAAGGTTGAATCGCTGGA | CGGCGATAGTCGTTAGCTTC |
| H2-T23 | GGACCGCGAATGACATAGC | GCACCTCAGGGTGACTTCAT |
| Serping1 | ACAGCCCCCTCTGAATTCTT | GGATGCTCTCCAAGTTGCTC |
| H2D1 | TCCGAGATTGTAAAGCGTGAAGA | ACAGGGCAGTGCAGGGATAG |
| Clcf1 | CTTCAATCCTCCTCGACTGG | TACGTCGGAGTTCAGCTGTG |
| Tgm1 | CTGTTGGTCCCGTCCCAAA | GGACCTTCCATTGTGCCTGG |
| S100a10 | CCTCTGGCTGTGGACAAAAT | CTGCTCACAAGAAGCAGTGG |
| Gapdh | CCAGAACATCATCCCTGCAT | TACTTGGCAGGTTTCTCCAG |
