## Supplementary figures and images for "A novel histone deacetylase inhibitor-based approach to eliminate microglia and retain astrocyte properties in glial cell culture"

### Supplemental Figure 1

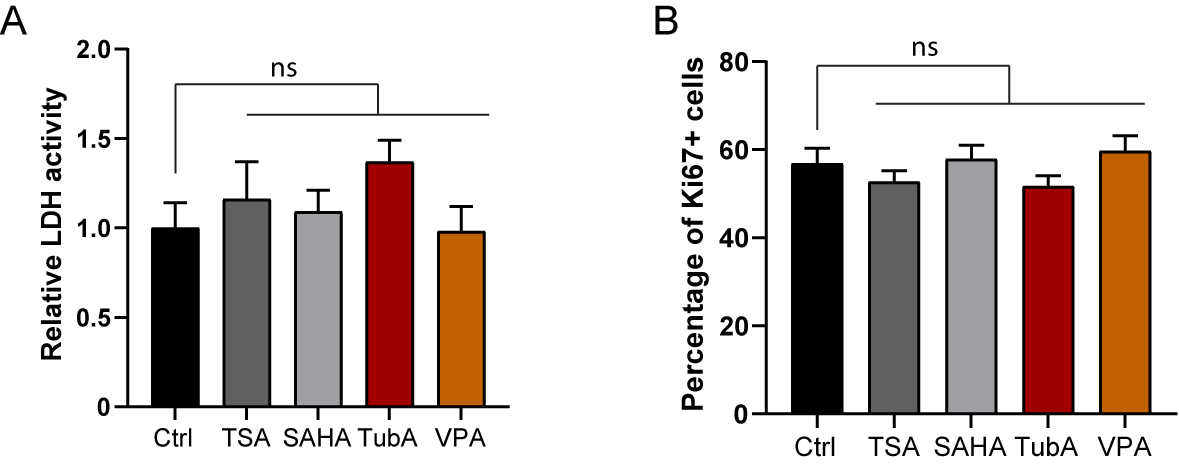

### Supplemental Figure 2

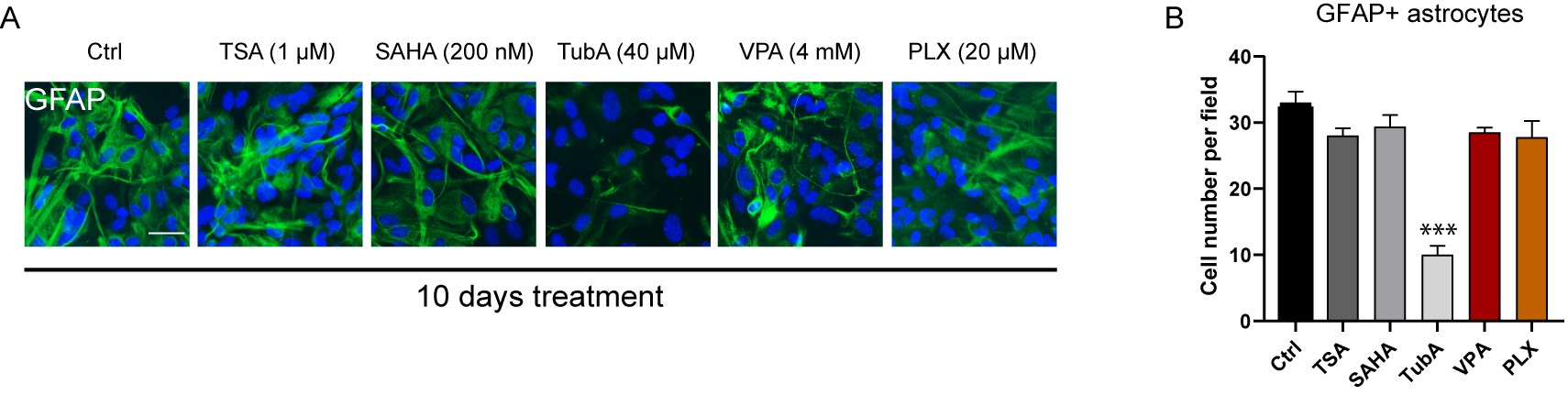

### Supplemental Figure 3

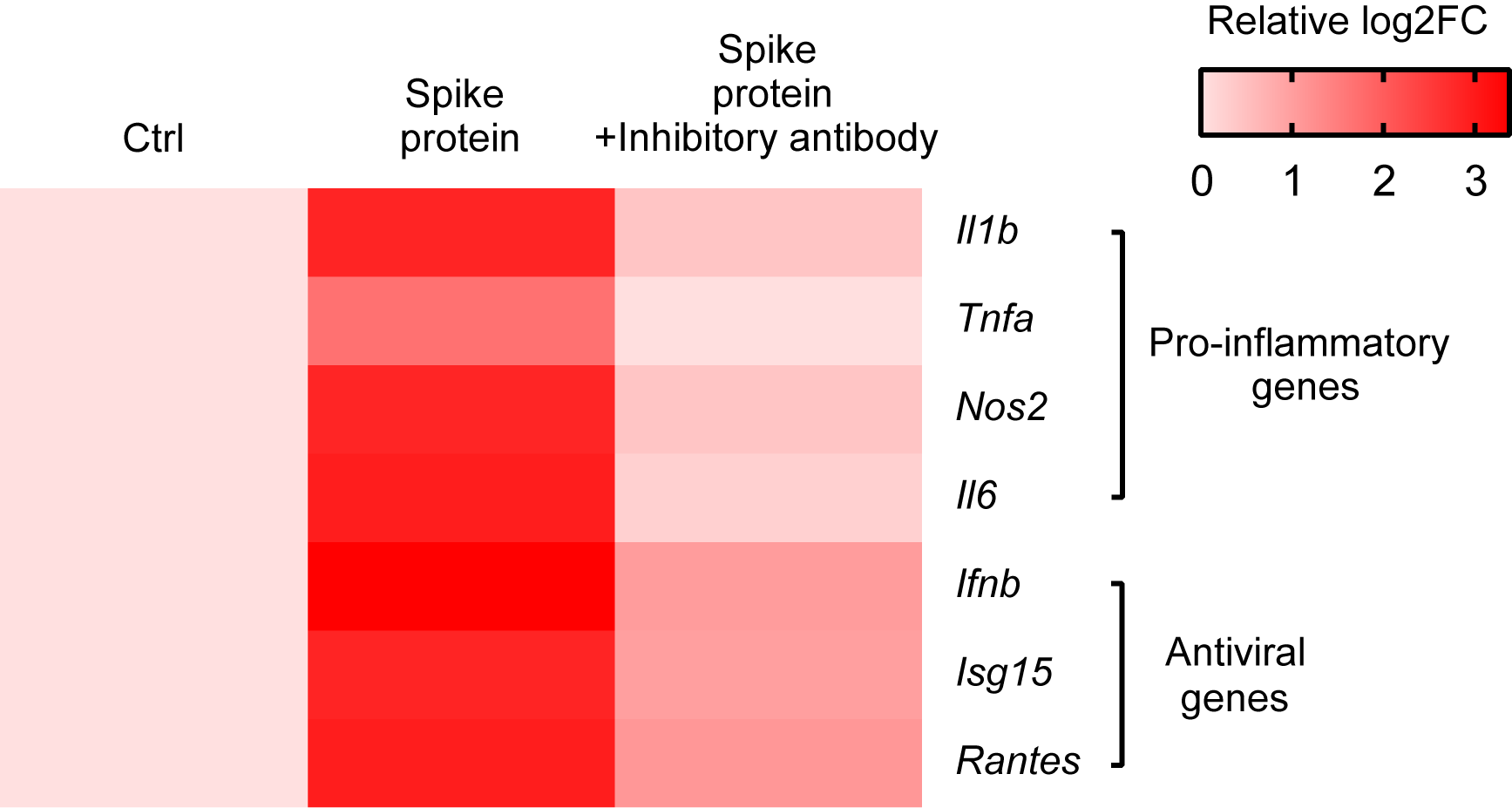
